# Enhanced proteome relative quantification using refined quantotypic spectral libraries

**DOI:** 10.64898/2026.07.06.736793

**Authors:** Bethany A. Barnes, Haneen Alharbi, Richard D. Unwin

## Abstract

Plasma proteomics is used for a variety of applications including biomarker discovery, disease monitoring, and drug development. Data-independent acquisition (DIA) has vastly improved the breadth of proteins that are identified from samples; however, given challenges in reproducibility and translation, it is critical that the quantitative performance of these methods is reliable.

Analysis of global proteomics data typically incorporates information from all detected peptides. However, some peptides do not reflect their parent protein amount, due to irreproducible digestion, modification, analytical interferences or instability. We hypothesise that including these peptides impacts protein relative quantification, and thus, a refined spectral library containing only quantitatively representative peptides provides superior protein quantification.

By analysing a defined multi-species spike-in model, we show that refining a plasma spectral library by removing precursors that fail to meet quality control metrics (25.4% of all identified precursors) reduces noise and variability, improving precision, accuracy and differential abundance analysis by up to ∼11%, with minimal identification losses and substantial reduction in computational demand. This demonstrates proof-of-concept that refining spectral libraries produces results that prioritize quantification quality over quantity. This approach could enable development of universal tissue-specific refined spectral libraries able to improve quantification quality with easy implementation and minimal processing time.

**Significance of the Study:** As DIA mass spectrometry proteome depth increases, the quality of the associated protein quantifications must be considered alongside identification breadth, particularly in complex matrices such as plasma, which presents additional technical challenges. The spectral library used for protein identification and quantification is a critical determinant of DIA performance, and its composition requires considerable consideration. This work illustrates an initial step toward improving protein quantification starting at the spectral library level by filtering precursors which are poor quantitative representatives of their parent proteins. In doing so, the resulting data is more reliable for downstream and biological interpretation, with fewer false differential abundance assignments and reduced quantitative noise. As such, this work represents a broader shift away from the habitual focus of MS workflows on maximising the number of protein and differential abundance identifications and instead prioritises the quality of quantification over quantity. These initial findings lay the groundwork for further development of spectral library refinement strategies, with the potential to continue improving the accuracy and precision of protein quantification in DIA-based proteomics.

## 1 Introduction

The human plasma proteome represents a vital source of readily available and clinically relevant data. As a diverse range of proteins are found circulating in plasma from various tissues throughout the body, plasma proteomics can provide a comprehensive overview of an individual’s health or disease status (Anderson and Anderson, 2002). Changes in plasma protein concentrations not only indicate disease and allow monitoring of disease progression but can also provide insight into biological processes which drive disease pathology (Birhanu, 2023). As such, plasma is an ideal biological sample for diagnostic applications as well as for biomarker, drug target and treatment discovery (Carrasco-Zanini et al., 2024).

Unfortunately, quantitative plasma proteomics has long been hampered by multiple challenges, including the heterogeneous, complex and dynamic nature of plasma (Anderson, 2004; Gerszten et al., 2008). Advances in mass spectrometry technology–particularly including the advent of Data Independent Acquisition (DIA) and the progression of associated data processing software – have advanced the ability to generate useful findings from plasma studies. DIA has improved plasma proteome coverage with increased sensitivity, accuracy and fewer missing values compared to data-dependent acquisition (Woo and Zhang, 2023; Metatla et al., 2024). DIA is still affected by some traditional mass spectrometry limitations, however. Specifically, while identification depth has vastly improved with recent algorithmic advances, uncertainty regarding the quality of label-free protein quantification remains. This is highlighted by Grossmann et al. (2026) in their paper introducing the novel quantification algorithm, QuantUMS. They draw attention to two persistent limitations: the difficulty of robustly quantifying proteins and peptides affected by high noise and signal interference, particularly as emerging machine learning approaches continue to expand their identifiability; and the challenge of estimating quantification errors in the absence of reliable quality control measures for peptide and protein abundances.

High-quality spectral libraries are a necessity for producing robust identification and quantification data in DIA-MS studies. Typically, peptides from a given proteome or specific sample are included in spectral libraries, either from *in silico* predictions from a protein database or from extant MS datasets, regardless of their performance quality. However, not all peptides perform equally. Consequently, the inclusion of quantitatively poor-quality peptides can affect overall protein quantification by causing biased intensity estimations, increased noise and errors, and even incorrect identification (Bogdanow, Zauber and Selbach, 2016; Dincer et al., 2022). Peptides considered poor for quantification occur due to sequence-specific biases impacting digestion, fragmentation, ionisation efficiency and chromatography behaviour as well as co-eluting interferences – all of which impact the ability of certain peptides to be reliably detected and accurately quantified by mass spectrometry (Abbatiello et al, 2010; Dincer et al., 2022; Liigand, Kaupmees and Kruve, 2019; Cappadona et al., 2012). This variability in quantitative reliability is seen clearly when optimising peptide panels for targeted measurements by Selected Reaction Monitoring, for example. Optimising DIA libraries for peptides considered suitable for inclusion in DIA-MS workflows, could limit these effects. A successful approach to peptide optimisation has been demonstrated with the TEAQ software which assesses the quality of peptides for inclusion in biomarker discovery by filtering peptides on a set of quality control measures including reproducibility and data completeness, effectively pre-validating candidate biomarkers (Fu et al, 2024).

Current popular quantification approaches, such as Top3, IBAQ or MaxLFQ, quantify all peptides from the spectral library which are identified in the data samples, then retrospectively assess these peptides for use in protein quantification using various criteria or thresholds such as quality scores or intensity levels, or include all peptide information in the final quantification (Cox et al., 2014; Silva et al., 2006; Schwanhausser et al., 2011). These methods have drawbacks regarding protein quantification accuracy, not limited to over- or underestimation of protein intensity and missing values, and also high computational requirements (Zhao et al., 2020; Krey et al., 2015).

QuantUMS is a novel quantification method implemented in more recent versions of the DIA processing software DIA-NN (from v.1.8.1 onward) which improves upon the above approaches by following a MaxLFQ-like algorithm while incorporating quantification uncertainty estimates taken at the individual signal level, as well as concordance and accuracy metrics for individual precursor and protein quantities. This enables effective data filtering to improve downstream statistical testing and interpretation. QuantUMS has been shown to improve accuracy and precision, to reduce bias, and improve differential abundance analysis (DAA) compared to other popular quantification methods (Demichev et al., 2020; Grossmann et al., 2026). However, whilst QuantUMS performs well, it is still likely to be prone to the effects of interferences, low signal, high noise, and both known and unknown peptide modification. As such we hypothesise that further progress can be made to improve the accuracy and precision of protein quantification and to reduce the demand on computational resources and processing time by optimising the spectral library pre-analysis rather than solely assessing peptide quality during (or post-) analysis.

Spectral library composition can have a substantial impact on abundance estimation. For example, Parker, Venkatraman and Van Eyk (2016) demonstrated surprisingly little overlap in statistically significant differentially abundant proteins observed when analysing the same dataset with three different spectral libraries. Previous studies have aimed to computationally optimise DIA spectral libraries through methods such as deep learning models (e.g. Carafe; Wen et al., 2025a), spectrum-centric conversion and analysis (MSLibrarian; Isaksson et al., 2021) or generating a smaller, experiment-specific subset library based on preliminary analysis (SubLib; Ge et al., 2021). These approaches have primarily focused on techniques to improve proteome coverage such as increasing peptide detection or accuracy of peptide identification. Spectral library optimisation with a direct focus on quantification robustness has not been duly explored.

Here we aim to refine protein quantification by generating a filtered spectral library containing only high-quality and quantitatively reliable peptides for analysis. As proof-of-concept, we have used mixed-species proteome samples with known, defined protein ratios; we have produced a ground truth dataset with known changes and constancy in protein abundance, enabling the filtering of poor-quality precursors and production of a new “mini” spectral library, consisting solely of peptides deemed suitable for quantification (Figure 1). When these data are reanalysed in DIA-NN, this so-called “mini library” reduces error rates and improves accuracy and precision of protein quantification, whilst retaining the majority of protein identifications. It is anticipated that this principle will allow the generation of universal, refined sample type-specific spectral libraries for easy and robust quantification analysis.

**Figure 1.**
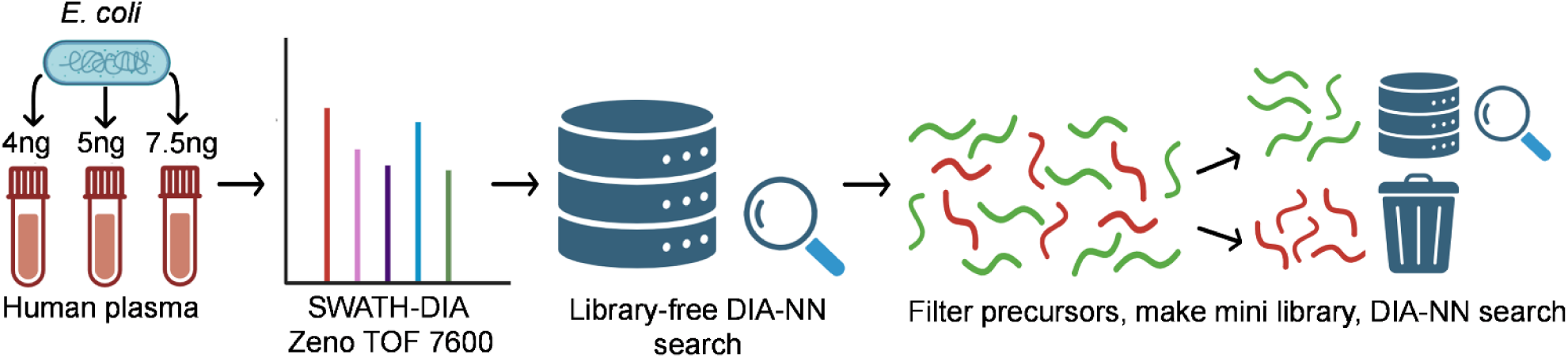
Schematic overview of the experimental design. Three concentrations of *E. coli* lysate were spiked into constant concentration depleted human plasma lysate and prepped for SWATH-DIA analysis. DIA data was analysed using an *in silico-*predicted library in DIA-NN (v.2.20). Identified precursors were filtered according to multiple quality measures to identify precursors with superior quantitative performance. A mini spectral library containing only these high-quality precursors was created, and the data was reanalysed in DIA-NN using this refined, “mini” library to yield improved quantification results.

## 2 Methods

### 2.1 Sample Preparation

Briefly, *E. coli* lysate was produced and spiked into 100 µg depleted (Top 14 Abundant Protein depletion, Thermo) human plasma (Sigma Aldrich) in 3 concentrations (4ng, 5ng and 7.5ng for the main dataset). Concentrations were determined by BCA protein assay (Thermo). Plasma-*E. coli* mixtures were digested using S-TRAP plates according to the manufacturer’s instructions (PROTIFI) and dried in a SpeedVac. Each concentration was produced and processed as full technical replicates (n=5), hereafter labelled E4, E5 and E7.5.

*E. coli* lysate spiked at concentrations of 0.6ng, 0.8ng and 1.2ng into 100 µg depleted human plasma with technical replicates n=5, was used as an additional dataset for testing using the same method as above.

### 2.2 LC-MS/MS Methods

Samples were reconstituted in a master mix of 20 iRT (index retention time) standards (JPT), 100 fmol/µl of PepCalMix (MS Synthetic Peptide Calibration Kit, SCIEX Ltd, UK), and loading buffer (2% (v/v) acetonitrile/0.1% (v/v) formic acid) at 0.16 µg/µl.

For the main dataset (*E. coli* concentrations spiked at 4ng, 5ng and 7.5ng), samples were injected at 560 ng/3.5 µl and analysed on the Zeno TOF 7600 (SCIEX) coupled with LC (Waters ACQUITY M-Class). The flow rate was set at 6 μL/min, with a gradient duration of 55 minutes. SWATH-MS was performed using 100 variable precursor windows. The TOF MS scan was acquired over the mass range (m/z) 400 to 1250. The total scan time was 1.543 s.

For the alternative sample set an optimised method with 65 variable windows was used. The TOF MS scan was acquired over the mass range (*m/z*) of 400 to 750. The total scan time was 1.171 s. The flow rate was set at 50 μL/min, with a gradient duration set to 21 minutes (27 minutes run time).

### 2.3 Data Analysis

DIA data was analysed by DIA-NN version 2.2.0 (Demichev et al., 2019). A predicted spectral library of human and *E. coli* proteins was generated by DIA-NN from a merged FASTA derived from published human and *E. coli* UniProtKB/Swiss-Prot reference proteomes (release 2022_01, proteome ID UP000005640; release 2022_01, proteome ID UP000000625). Five .wiff technical replicate output files of each *E. coli* concentration (E4, E5 and E7.5) were converted to .dia, and Mass accuracy was set to 20.0 and MS1 accuracy to 12.0 as recommended by the developer. Match between runs (MBR) and protein inference were enabled, threads were set to 14 and all other settings left as default.

### 2.4 Mini library generation

*E. coli* spiked human plasma was analysed by mass spectrometry and DIA-NN (v2.2.0) was used for protein identification and quantification as described above. The *in silico-*predicted spectral library was filtered to produce a mini library by importing the precursor matrix output file from DIA-NN into R and calculating a set of quality metrics for each identified precursor. These quality metrics included: the mean of the 5 technical replicates for each of E4, E5 and E7.5 concentrations, the coefficient of variation (CV) of E4, E5, and E7.5 samples, the ratio of the mean intensity of E4, E5, and E7.5 relative to the E4 mean, Pearson correlation coefficient (R) and coefficient of determination (R²) between the spike-in ratio, 1, 1.25, 1.875 (4ng/4ng, 5ng/4ng and 7.5ng/4ng), and the aforementioned mean intensity ratio and the number of missing values per precursor across all replicate samples.

Multiple combinations and cut-offs were tested for each of these quality metrics (see Supplementary Table 1). For the final mini library, precursors were retained if they met the following optimal species-specific parameters. Human precursors were retained if they exhibited CV <30% in at least 2 of 3 concentrations, identification in ≥3 of 5 technical replicates for each spike-in concentration, and either R² <0.4 or mean intensity ratios within ±12% of expected values. *E. coli* precursors were retained if they exhibited CV <30% in at least 2 of 3 concentrations, identification in ≥3 of 5 technical replicates for each spike-in concentration, and either R >0 and R² >0.6 or mean intensity ratios within ±12% of expected values. For only those precursors deemed good quality, the peptide sequence and associated spectral library data for the precursor and all associated features was incorporated into the mini spectral library.

All samples were reanalysed in DIA-NN using the new mini spectral library and otherwise identical parameters to generate refined quantification results.

#### 2.4.1 Reanalysis with Refined Libraries

Two types of data reanalyses were explored. The first, which we term the “empirical library analysis” used the empirical spectral library generated from the second-pass of the initial DIA-NN analysis. The second type of reanalysis used our refined “mini library” containing only precursors which matched the quality filters described above. In both reanalyses, parameters and settings remained identical to the original “predicted library” analysis, differing only in the spectral library applied.

### 2.5 Statistical Analysis

#### 2.5.1 Binary classification

Protein group comparisons between the E4 and E7.5 samples were performed using two-tailed Welch’s t-tests under multiple analytical conditions: complete-case analysis (identifications with missing values excluded); mean imputation, whereby protein groups with fewer than three missing values had these replaced with the mean intensity for that protein, while those with three or more missing values were excluded; both approaches with identifications having one or fewer proteotypic peptides excluded; and analysis permitting unequal sample sizes. True positives were defined as *E. coli* proteins with a statistically significant p-value, true negatives as human proteins with a non-significant p-value, false positives as human proteins with a significant p-value, and false negatives as *E. coli* proteins with a non-significant p-value.

#### 2.5.2 Linear regression modelling

A logistic regression model was fitted using the p-value from the Welch’s t-test between E4 and E7.5 to estimate the probability of correct species categorisation using the glm() function in R with a binomial family specification. To train the model, 70% of the data was randomly selected and the remaining 30% was used as testing data. This model, along with the R package pROC (Robin et al., 2011), was used to generate receiver operating characteristic (ROC) curves and calculate area under the curve (AUC) values.

#### 2.5.3 Differential Abundance Analysis (DAA)

For DAA, log_2_ fold changes (log_2_FC) were calculated between each pairwise comparison and the associated p-value from the Welch’s t-test was adjusted using the Benjamini-Hochberg method. Proteins were considered significantly differentially abundant if the 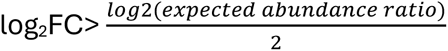 and the adjusted p-value <0.05.

## 3 Results

The primary aim of this study was to develop a refined, dataset-specific spectral library which could improve quantification quality when processing data with the software tool DIA-NN. We generated a ground-truth dataset using DIA-MS analysis of *E. coli* lysate spiked at three concentrations (4ng, 5ng and 7.5ng) into a human plasma background maintained at a constant concentration. This design provided a benchmark of expected results for evaluation of binary classifications, protein abundance ratios and differential abundance.

Following initial DIA-NN analysis, precursors were filtered against several quality criteria including CV, R^2^ and intensity ratios. Precursors meeting these criteria were deemed quantitively representative of their parent protein and compiled with their corresponding spectral information to generate a “mini” spectral library. We iteratively tested 53 filtering parameter combinations, evaluating each method based on the number of identifications, the binary classification performance, CV, number of proteotypic sequences and missingness (Supplementary Figure 1). We optimised the filtering methods, prioritising maximisation of true positives and maintenance of low false classification rates, with as few protein identification losses as possible (see Supplementary Table 1). The chosen filtering method required retained human precursors to have a CV <30%, identification in ≥3 of 5 technical replicates and either R² <0.4 or intensity ratios within ±12% of expected values. Similarly, *E. coli* precursors were required to have a CV <30% in at least 2 of 3 concentrations, identification in ≥3 of 5 technical replicates, but then either R >0 and R² >0.6 or intensity ratios within ±12% of expected values. Of the 9,625 unique precursors identified within the initial analysis, 2,449 (25.4%) were deemed poor quality and were excluded from the spectral library. Precursors which were not identified in the samples were also excluded from the mini library – approximately 4.8million.

This generated a refined, “mini” library of 840 protein groups and 7,167 precursors which was used to reanalyse the data in DIA-NN. By excluding unidentified and poorly performing precursors from both the spectral library and subsequent protein quantification, we hypothesized that the resulting quantification would be less affected by noise, as well as exhibiting improved accuracy and enhanced precision. A schematic overview of this workflow is shown in Figure 1.

For comparison, the data was also reanalysed using the second-pass empirical spectral library generated by DIA-NN (referred to here as the “empirical” library) due to its smaller size (1,211 protein groups and 9,882 precursors) with increased relevance to the specific datasets being analysed. According to the DIA-NN developers, this second-pass library can improve identification rates, data completeness, and quantification accuracy (https://github.com/vdemichev/DiaNN).

### 3.1 Filtering spectral library precursors can maintain protein identification rates whilst reducing computational demand

Following DIA-NN analysis of the SWATH data using the *in silico*-predicted, empirical, and mini spectral libraries, we first evaluated the protein identification performance of each library. Despite the vast reduction in library size of the empirical and mini libraries, with both containing substantially fewer precursors and proteins than the *in silico*-predicted library, the number of peptide and protein identifications decreased only moderately: 797, 756, and 707 protein identifications were obtained from the predicted, empirical, and mini library, respectively (Figure 2A). Additionally, the majority of protein identifications were shared by all three spectral libraries (Figure 2B), with ∼85% of the 59 identifications found only in one library search lacking proteotypic peptides.

**Figure 2.**
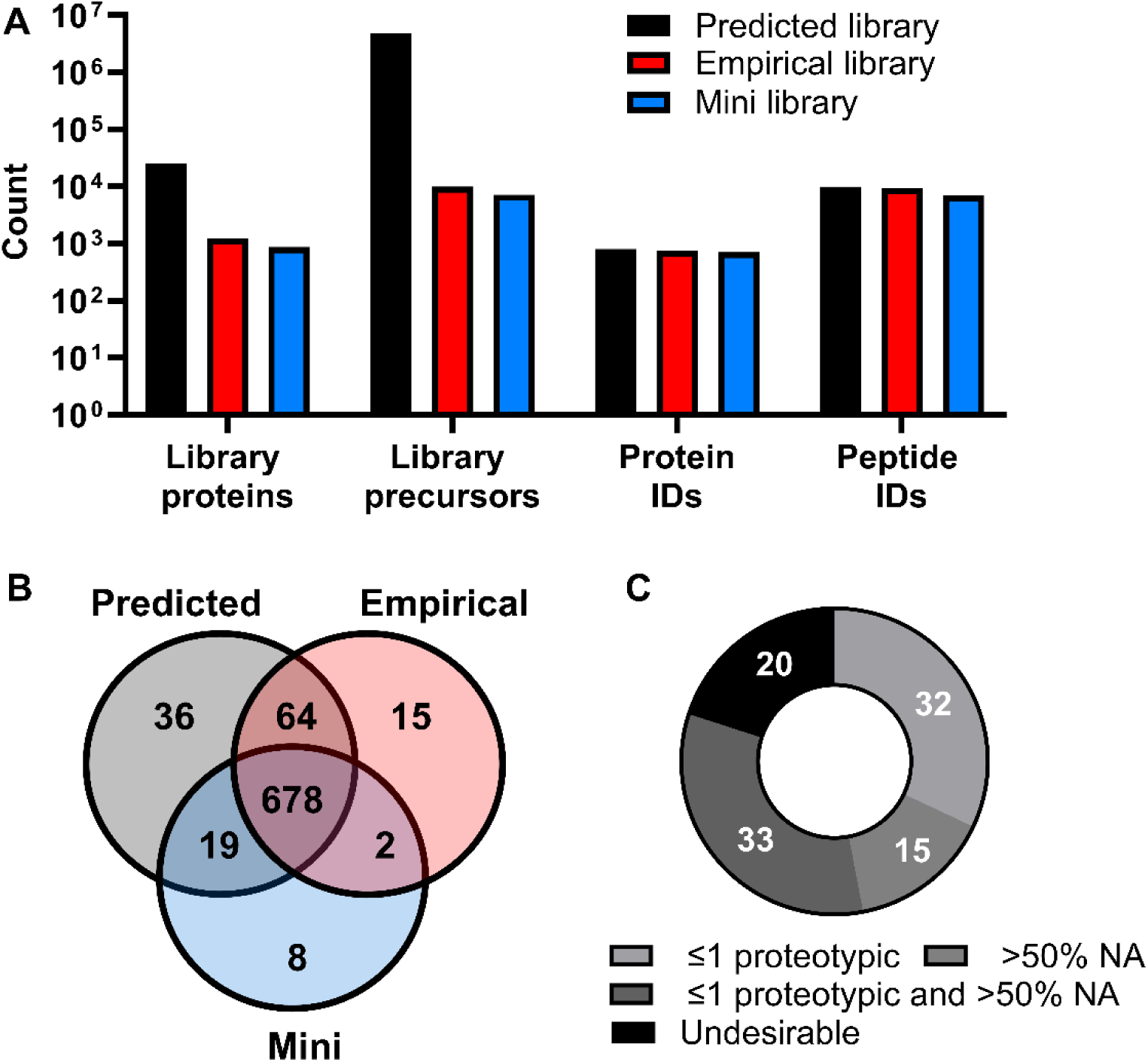
Mini library retains bulk of protein IDs. (A) Library size and corresponding identification rates across the three spectral libraries. (B) Overlap of protein identifications between libraries shown by Venn diagram. (C) Proteins identified in the *in silico* predicted library but lost in the mini library, categorized by quality metrics: proteins with ≤1 proteotypic peptide, proteins with >50% missing values (NA), proteins meeting both criteria, and proteins classified as “undesirable” losses (>1 proteotypic peptide and <50% missing values).

To assess the identifications that were lost upon re-analysis with the mini library, we examined the number of proteotypic peptides and missing values for these proteins. Proteins with one or fewer proteotypic peptides have a higher risk of false-positive identification, unreliable quantification, and greater technical variability (Carr et al., 2004). Similarly, missing values reduce statistical power and can introduce substantial bias, resulting in inaccurate protein measurements (Bramer et al., 2020; Harris et al., 2023). Both metrics were therefore used as quality indicators.

Approximately 80% of “lost” identifications had either one or no proteotypic peptides, more than half missing values, or both. Due to the inherent problems associated with quantification based on non-proteotypic peptides or extensive missing values, the quantification of these identifications likely exhibited poor quality or lacked reliability, thus, we considered their loss desirable. A relatively small proportion of lost identifications (n=20) had more than one proteotypic peptide and fewer than half missing values (Figure 2C). Further investigation of these proteins revealed that in 16 of 20 cases, no precursors met the high-quality threshold and were therefore absent from the mini library. The remaining four proteins had one precursor each retained in the mini library, but were not re-identified in the data; however, these four proteins represented only 0.5% of the total protein identifications from the predicted library.

The large reduction in spectral library size – from an approximate 9.6 GB for the predicted library to 22 MB (∼99.8% reduction) for the mini library in a .tsv format – dramatically reduced computational demand and processing time, decreasing analysis time per datafile from over one hour for 15 samples to approximately 6 minutes. This is reported whilst acknowledging that these timings are machine-dependent and reflect the processing power of the hardware used. Nevertheless, the efficiency gains were substantial. Although the initial analysis for mini library generation required significant time and computational resources, the mini library approach offers clear advantages for repeated analyses and could prove particularly valuable for large cohort studies.

### 3.2 Mini library enhances correct binary classification

To compare the efficacy of the predicted, empirical and mini spectral libraries when used for DIA-NN analysis, we evaluated the effect of each library on the classification of proteins based on species as a surrogate for either altered or unchanged abundance. A two-tailed Welch’s t-test was performed between the five replicates containing 4ng of *E. coli* spike-in (E4) and five replicates containing 7.5ng *E. coli* spike-in (E7.5), with statistical significance set at p<0.05. True positives were deemed to be *E. coli* proteins with significant differences between groups, and true negatives as human proteins exhibiting no significant change.

Proteins with missing quantification values in any replicate were initially excluded from the t-test to ensure equal sample sizes, maintaining statistical power and avoiding sensitivity to unequal variances. When comparing the predicted and mini libraries, we observed a modest reduction in true negatives (TN) from 283 to 275 and a small increase in false negatives (FN) from 2 to 5. However, true positives (TP) increased substantially from 281 to 313, while false positives (FP) decreased by nearly two-thirds (Figure 3A). Additionally, fewer missing values were present in the mini library compared to both the empirical and predicted libraries. (Figure 3B).

**Figure 3.**
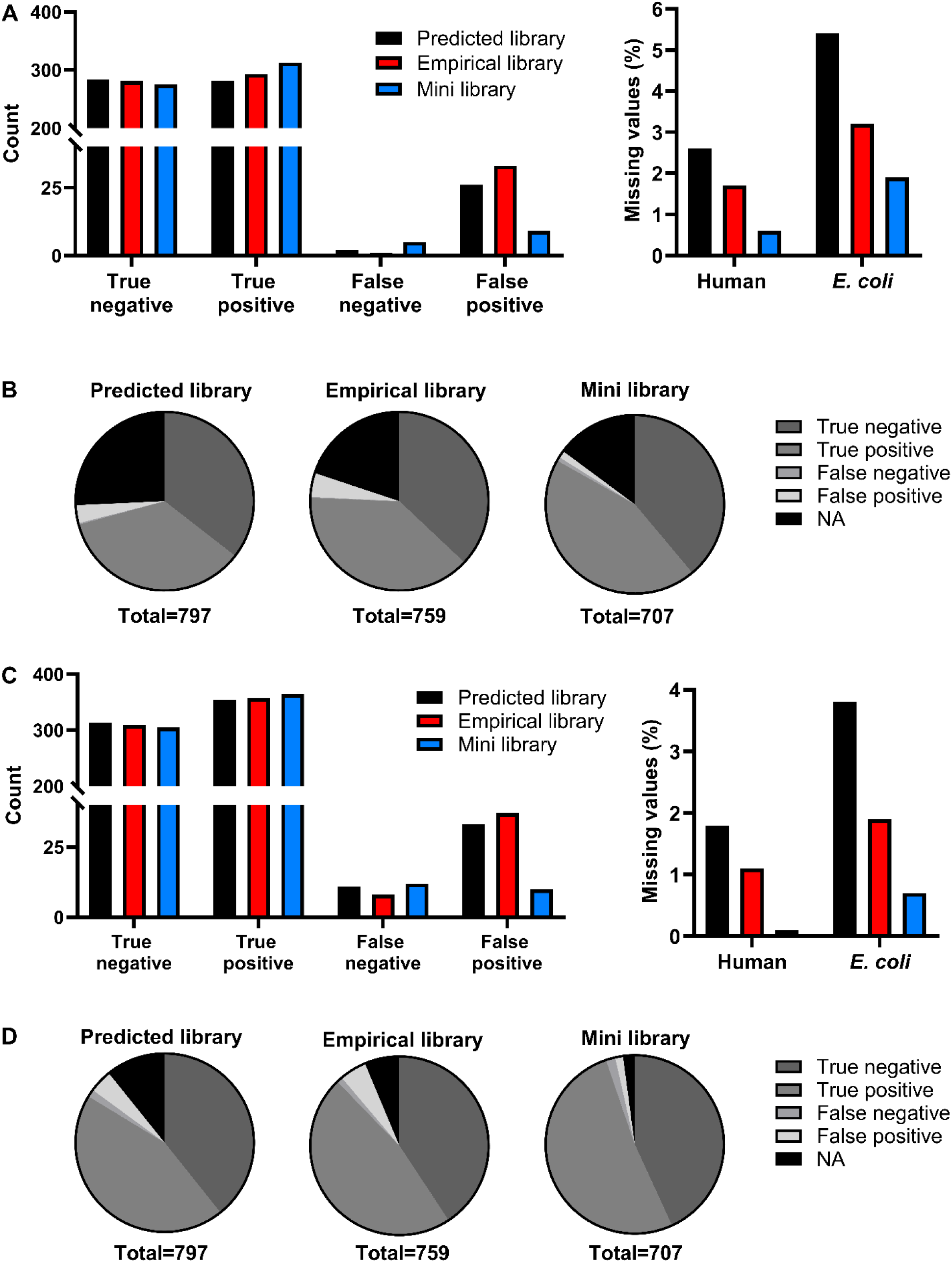
Mini library increases correct binary classification across multiple analytical approaches. An unpaired Student’s t-test was performed between E4 and E7.5 protein quantifications for human and *E. coli* identifications to determine true positives (TP; significantly changed *E. coli* proteins), true negatives (TN; stable human proteins), false positives (FP; falsely significant human proteins), and false negatives (FN; non-significant *E. coli* proteins). (A) Raw counts of TP, TN, FP, and FN classifications for each library using only proteins with complete data across all five technical replicates. (B) Total number of missing values (NA) across technical replicates for all protein identifications with each spectral library. (C) Proportions of TP, TN, FP, FN, and NA (excluded identifications) categories from panel A for each library. (D) Raw counts of TP, TN, FP, FN, and NA classifications for each library using mean imputation for proteins with fewer than three missing values; proteins with three or more missing values were excluded. (E) Total number of missing values (NA) across technical replicates for all protein identifications with each spectral library after mean imputation for eligible proteins. (F) Proportions of TP, TN, FP, FN, and NA categories from panel D for each library.

Although we observed a reduction in the count of true negatives with the mini library, when expressed as a proportion of total protein identifications correct TN classification actually increased from 35.5% with the predicted library to 38.9% with the mini library. This suggested that the reduction in the number of true negatives reflected overall protein identification loss rather than reduced accuracy of classification. Similarly, true positives increased from 35.3% to 44.3% proportionally, while false positives decreased from 3.3% to 1.3% and the proportion of excluded identifications also decreased. Despite a modest increase in FN count, proportionately false negatives represented only approximately 0.7% of identifications with the mini library compared to 0.25% with the predicted library, indicating minimal practical impact. (Figure 3C).

We repeated the analysis using mean imputation for proteins with up to two missing values, excluding any proteins with more than two missing values. This approach provided a more complete classification overview including most identified proteins. The results mirrored the complete-case analysis: a slight reduction in TN counts due to identification loss, increased TP, one additional FN, and a greater than two-thirds reduction in FP with the mini library (Figure 3D). Proportionally, the mini library approach outperformed the standard predicted library across true negatives, true positives, and false positives and the overall accuracy 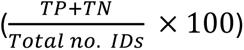 of the classification increased from 83.7% with the predicted library to 94.8% with the mini library (Figure 3F).

Similar increases in classification accuracy were acquired (89.3% overall accuracy with the predicted library vs 94.8% with the mini library) when identifications with one or no proteotypic peptides were removed – a standard practice in mass spectrometry data analysis due to the inherent unreliability of single-peptide identifications (Supplementary Figures 2A and 2B). Consistent outcomes were also observed when the t-test was conducted with unequal sample sizes including overall accuracy increasing from 81.3% with the predicted library to 91.5% with the mini library, although a larger decrease in true negative counts was evident under these conditions (Supplementary Figures 2C and 2D).

In general, the empirical library generated from the primary DIA-NN search demonstrated intermediate performance between the predicted and mini libraries, with fewer identification losses but smaller TP gains compared to the mini library, and persistent FP increases relative to both libraries (Figure 3A-F).

Thus, the mini library consistently improved the numbers of true positives and false positives, regardless of how missing values or single-peptide identifications were handled. While this came at the cost of modest identification losses, proportional performance measurements demonstrated that the mini library consistently outperformed both the predicted and empirical libraries across nearly all classification categories and analytical approaches (Supplementary Table 2).

### 3.3 Mini library produces data-set specific improvement of quantification quality

To further investigate the extent of improvement produced by the mini library approach we examined multiple binary classification metrics and standards of statistical reliability for the mean imputed classification set. We first assessed precision and recall at the protein level for both the predicted and mini library by tallying binary class labels for a range of p-value thresholds (0.0001 to 1.0) and calculating the corresponding precision and recall values. The mini library precision-recall curve remained above that of the predicted library across all thresholds (Figure 4A). This effectively illustrates the robustness of the binary classification using the mini library approach in this model across varying stringency thresholds.

**Figure 4.**
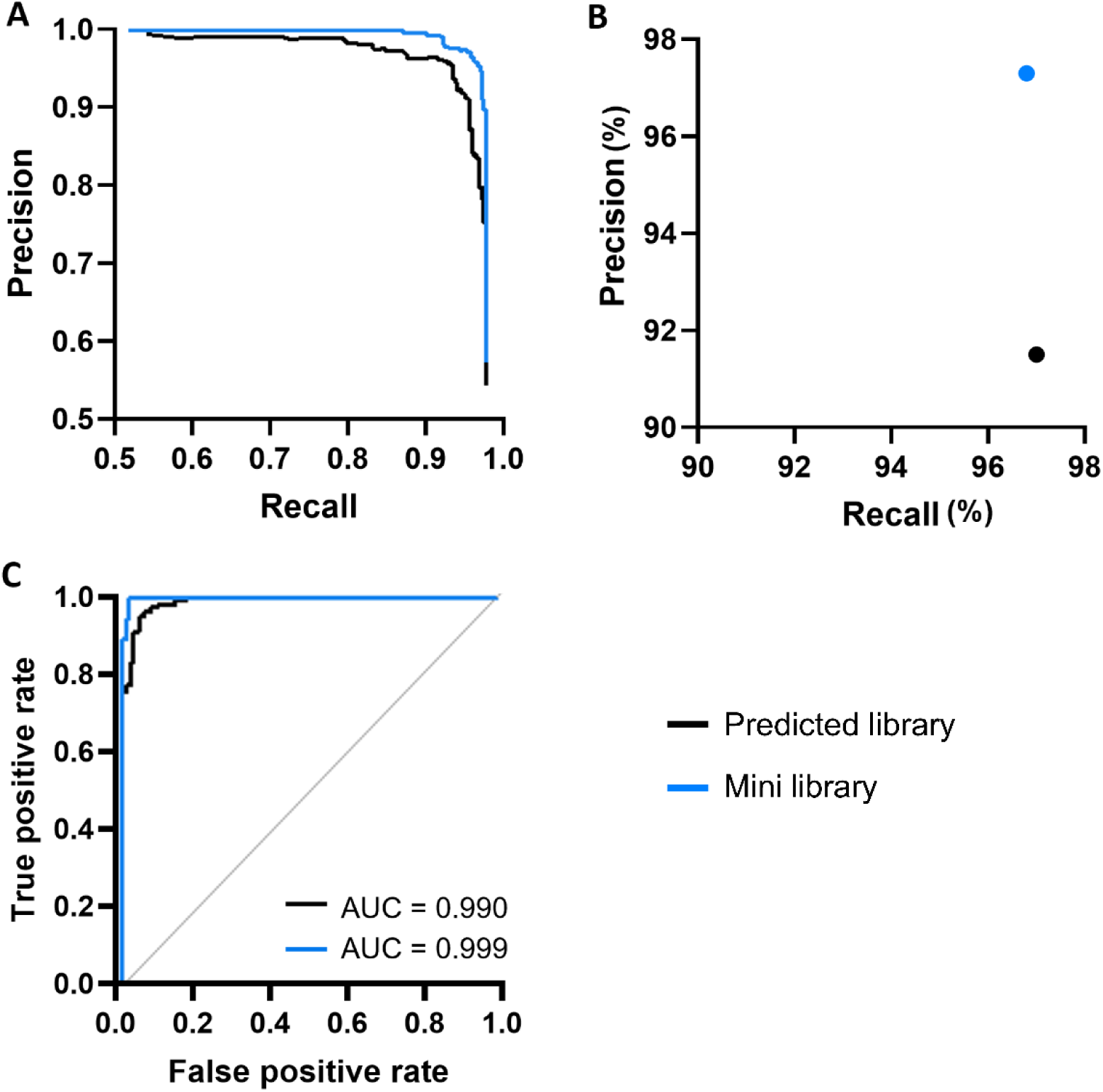
Improved recall with mini library application. Binary classification was evaluated by comparing precursor identification statistics against ground truth data. Performance was assessed using (A) Precision-recall curves across a range of p-value thresholds (0.0001 to 1.0) for both the predicted and mini library methods. (B) Precision and recall of the binary classifications at the standard significance threshold, p=0.05. (C) ROC curves and corresponding AUC values for the logistic regression model of p-value as a predictor of species, with a probability threshold of 0.7 applied to both library results.

We also evaluated precision and recall derived directly from the confusion matrix for this specific dataset with a p-value threshold of 0.05 (Figure 4B). A substantial increase in precision of approximately 6% was apparent with the mini library, with only a minimal associated decrease in recall, less than 0.5%. The mini library also demonstrated superior performance when assessed against additional quality metrics: false positive rate (FPR), accuracy, specificity and F1 score (Table 1). These results demonstrated that the modest loss in recall was an acceptable trade-off for generating a higher-quality dataset with improved classification performance.

**Table 1.**
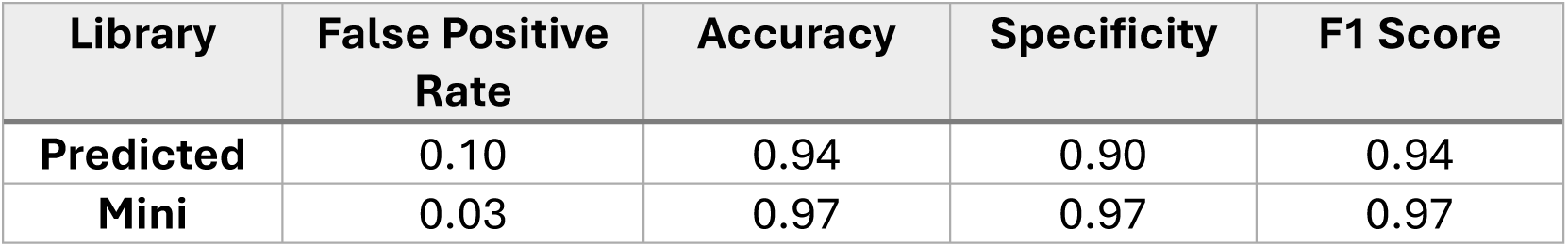
Mini library improves several standards of statistical reliability.

| Library | False Positive Rate | Accuracy | Specificity | F1 Score |
| --- | --- | --- | --- | --- |
| **Predicted** | 0.10 | 0.94 | 0.90 | 0.94 |
| **Mini** | 0.03 | 0.97 | 0.97 | 0.97 |

The p-values generated from the Welch’s t-test for the comparison between E4 and E7.5 were then employed to train a linear regression machine learning algorithm with a probability threshold of 0.5. This model was developed to evaluate the ability of each library to correctly predict species (as a surrogate for differential abundance) based upon p-value and to assess their potential applicability to larger or alternative datasets. Figure 4C presents ROC curves for the predicted and mini libraries alongside their corresponding AUC values. Both libraries demonstrated excellent discriminative performance, with an AUC of 0.990 (95% CI: 0.981–0.998, DeLong method) for the predicted library and an AUC of 0.999 (95% CI: 0.996–1.000, DeLong method) for the mini library. While the difference between the two curves was not statistically significant (*p* = 0.050), ROC/ AUC analysis slightly favoured the mini library, and the mini library performance excelled at higher levels of true positives and lower levels of false negatives demonstrating that the p-values generated when using the mini library for data analysis better reflected correct species classification and therefore correct identification of differentially abundant proteins.

### 3.4 Improved binary classification translates to enhanced quantification precision and accuracy

We next investigated whether the improved classification performance of the mini library translated to increases in the accuracy and precision of the protein abundance ratio between sample groups. LFQbench (Navarro et al., 2016) analysis demonstrated that the mini library substantially improved precision, resulting in markedly lower log-standard deviations for both precursor and protein quantity ratios. With the predicted library, human protein log_2_ratios were scattered with a standard deviation of 0.21 around the expected ratio, whereas this tightened to 0.17 with the mini library. Similarly, for *E. coli* proteins, standard deviation reduced from 0.26 with the predicted library to 0.21 with the mini library (Supplementary Table 3). This is an approximate 19% decrease in deviation for both species, demonstrating that fewer proteins are quantified erratically and more cluster consistently around the expected ratio with the mini library. The greatest improvements in precision were observed for low-intensity precursors (Supplementary Figure 3).

Accuracy of the protein abundance ratio (defined here as median deviation of log-ratios from the expected ratio) remained unchanged – median ratios in the predicted library closely matched expected values and no improvement was seen with the mini library. Encouragingly, the mini library exhibited lower technical variance (the median CV for the background species among replicate runs) of 6% vs. 8% for the mini library and predicted library, respectively, and decreased the overlap between species quantity ratios with the AUC of ROC curves assessing species separation ability of 0.999 for the mini library vs. 0.885 for the predicted library (Supplementary Table 4).

The improvement in precision at the peptide level appears to stem from the mini library filtering process effectively removing peptides where the intensity ratio deviates largely from expected values (Supplementary Figure 3A). When carried through to the protein level, this supports the hypothesis that this approach decreases identification of imprecisely quantified proteins. Visual analysis of the log-ratio scatter plots produced by LFQbench suggested that the mini library approach led to the undesirable loss of a small number of seemingly accurate proteins, particularly human proteins, and thus demonstrates room for improvement. However, this is weighed in balance with the mini library’s efficacy at removing low-quality proteins, and consequently noise from subsequent analysis (Supplementary Figure 3B).

We additionally hypothesised that the exclusion of poor-quality peptides would also result in improved quantification accuracy of retained proteins. To explore this further we compared the proportion of proteins with mean abundance ratios falling within ±20% of expected abundance ratios for each library. Each *E. coli* spike-in replicate set (E4, E5, and E7.5) was compared pairwise, producing expected abundance ratios of 1.875× for E7.5 vs. E4, 1.5× for E7.5 vs. E5, and 1.25× for E5 vs. E4. The proportion of quantifications falling within ±20% of these expected values was consistently higher for the mini library across all comparisons, with a mean improvement of 6.1% (Table 2).

**Table 2.** The count and proportion () of proteins with mean abundance ratios which lie within +/- 20% of the expected abundance ratio in the predicted and mini libraries for each concentration comparison.

| Ratio comparison | Human |  | *E. coli* |  |
| --- | --- | --- | --- | --- |
|  | Predicted | Mini | Predicted | Mini |
| **E7.5 vs E4** | 320<br>(86.7%) | 304<br>(95.9%) | 316<br>(73.8%) | 310<br>(79.5%) |
| **E7.5 vs E5** | 323<br>(87.5%) | 292<br>(92.1%) | 333<br>(77.8%) | 330<br>(84.6%) |
| **E5 vs E4** | 323<br>(87.5%) | 284<br>(89.6%) | 336<br>(78.5%) | 339<br>(86.9%) |

However, this improvement was not evident in the total count of proteins with abundance ratios lying within expected values. To determine whether this was simply an artefact of lower identification numbers with the mini library, we assessed only protein identifications that intersected between the two libraries as a direct and equal comparison of library performance – a total of 697 overlapping proteins. For proteins identified with both libraries, we observed a total increase of 6 proteins falling within ±20% of the expected values with the mini library for both the E7.5 vs E4 and E7.5 vs E5 comparisons. A slight decrease in the total number of proteins falling within ±20% of the expected values was seen (5 proteins) for the E5 vs E4 comparison due to decreased accuracy of human proteins (Supplementary Table 5).

These results show that the mini library improves the accuracy of protein relative abundances, although with a slightly more pronounced improvement for differentially abundant proteins.

### 3.5 Improved performance of mini library boosts differential abundance analysis (DAA)

The analysis above focused on binary classification based solely on a single statistical test, as the small fold changes in our model limit the discriminative power of fold-change-based filtering, causing statistical testing to give the more correct outcome. However, most studies commonly also employ a cut-off based on fold-change. As such, we investigated whether the above discussed improvements translated to enhanced performance in what we define a DAA, where the inclusion of a log_2_FC cut off prevents proteins with small fold-changes being considered significant.

We performed DAA, calculating log_2_FC between pairwise comparisons and adjusting the associated p-values from the Welch’s t-test using the Benjamini-Hochberg method. For the E4 vs E7.5 comparison, proteins were considered significantly differentially abundant if log_2_FC>0.45 and adjusted p-value <0.05. At low p-values (p<0.01), both libraries identified similar numbers of differentially abundant proteins (DAPs). However, the mini library demonstrated improved DAP identification beginning at p=0.01 and maintained this advantage up to p=0.1 (Figure 5A). When examining true DAPs for the same comparison the mini library correctly identified more DAPs (345 vs. 341) despite having fewer overall protein identifications. It almost halved the misclassification of *E. coli* proteins (45 vs 87) (Figure 5B). Proportionally, the combined accuracy for true DAPs and true stable proteins improved from 89.0% with the predicted library to 93.6% with the mini library (Figure 5C). The mini library also reduced false DAPs (0 vs. 1). While it identified fewer true stable proteins overall (317 vs. 368), this again appeared to reflect identification losses rather than decreased accuracy.

**Figure 5.**
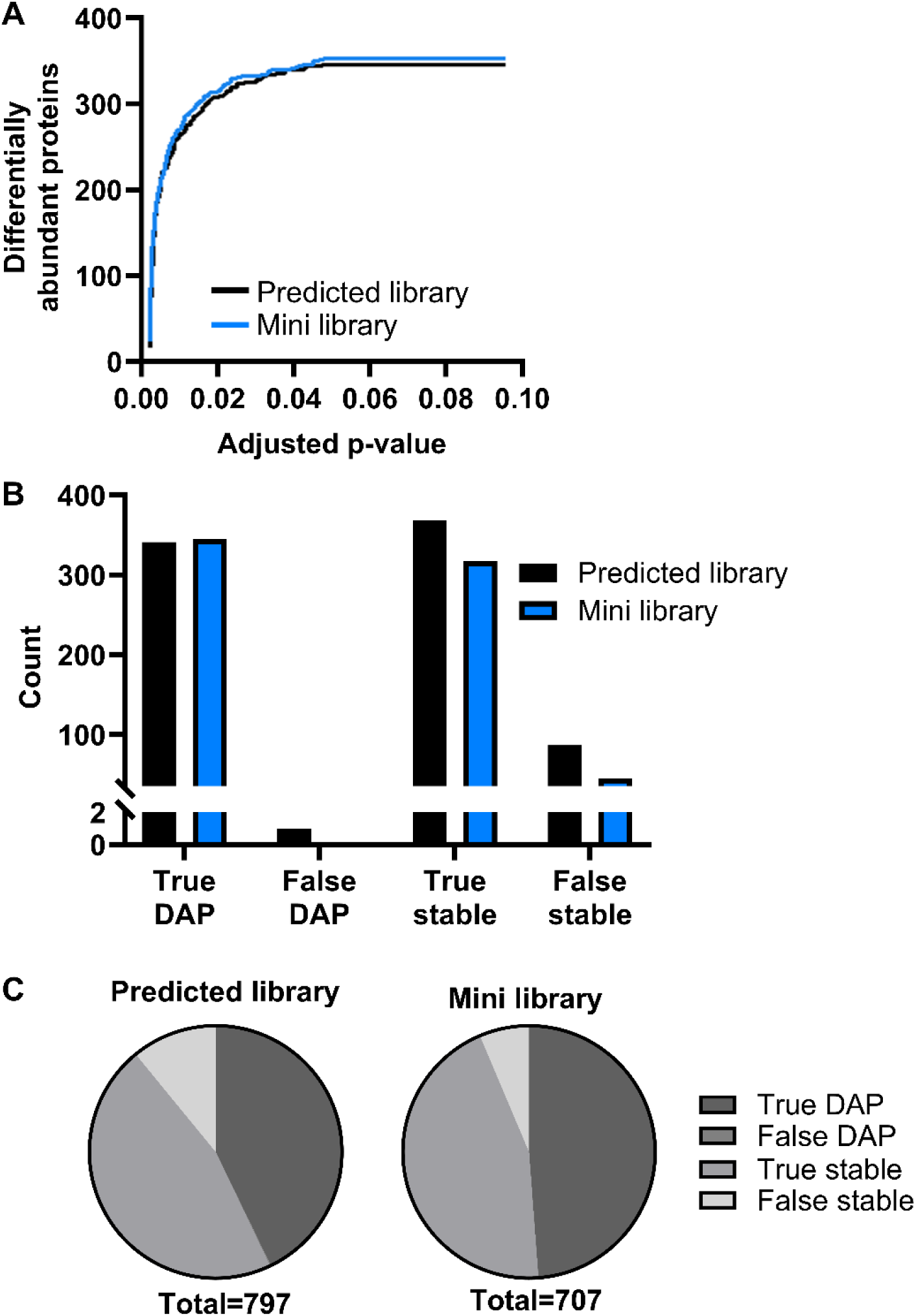
Mini library output outperforms the predicted library in differential abundance analysis. For the E4 vs E7.5 pairwise comparison (A) The number of differentially abundant proteins at ranging adjusted p-values from 0.001 to 0.1 for both the predicted and mini libraries. (B) The number of true differentially abundant proteins (true DAP; *E. coli* proteins with log_2_FC >0.45 and adjusted p-value <0.05), false differentially abundant proteins (false DAP; human proteins with log_2_FC >0.45 and adjusted p-value <0.05), true stable proteins (human proteins with log_2_FC <0.45 and/ or adjusted p-value >0.05) and false stable proteins (*E. coli* proteins with log_2_FC <0.45 and/ or adjusted p-value >0.05) for predicted and mini libraries. (C) The proportion of differentially abundant and stable proteins for each library.

To verify this, we examined only proteins identified by both libraries (Table 3). Again, differential abundance for the E4 vs. E7.5 comparison was defined as a log_2_FC >0.45 and an adjusted p-value <0.05. Among these intersecting proteins, the mini library identified more true DAPs (345 vs. 340) and fewer false stable proteins (45 vs. 50). Notably, neither library produced false DAPs among the intersecting proteins, and both identified the same number of true stable proteins. This confirmed that the reduction in true stable protein counts resulted entirely from identification loss rather than decreased DAA accuracy and established that the mini library DAA of *E. coli* proteins outperformed the predicted library and DAA of human proteins performed with at least the same accuracy as the predicted library. When considered alongside the improved true DAP identification and reduced false DAPs, the mini library demonstrated greater overall DAA accuracy for this pairwise comparison.

**Table 3.**
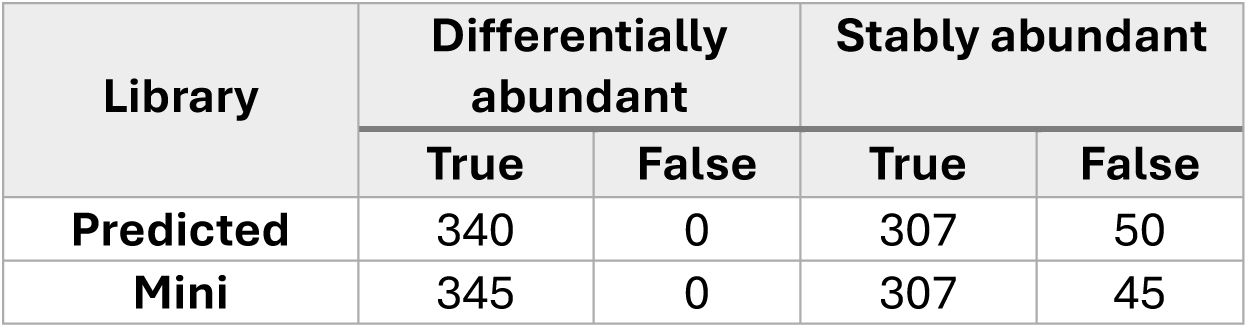
Number of differentially abundant and stable proteins in the pairwise comparison between E4 and E7.5, only from proteins identified by both libraries. Differentially abundant proteins are classed as those with a log_2_FC >0.45 and adjusted p-value <0.05.

Other comparisons can be seen in Supplementary Table 6 although the lower FC in the spike-in design makes these analyses less informative.

### 3.6 Application to alternative datasets

To test the generalisability of our mini library, or mini library generation workflow, to different datasets, we applied the mini library to the analysis of an alternative, independent *E. coli* spiked human plasma dataset. These data were still based on *E. coli* proteins spiked into plasma, maintaining the identities of the differentially abundant proteins and the matrix, but here the samples contained lower spike concentrations and different dilution factors (0.6ng, 0.8ng, and 1.2ng) compared to the original dataset, and were acquired across a shorter LC gradient for SWATH-MS, with different SWATH window settings. The alternative dataset was also run with the *in silico*-predicted library to generate a new performance baseline to compare against.

When using the mini spectral library generated from the first dataset for DIA-NN analysis with this alternative dataset, the number of overall identifications increased from 397 with the predicted library to 402 with the mini library, as did the number and proportion of true positive quantifications, 122 to 131 (30.7% to 32.6%). All other classification metrics (TN, 240 vs 218; FP, 19 vs 22; and FN, 16 vs 31) worsened with the mini library approach (Supplementary Figure 4), as did all other statistical quality metrics.

In contrast, using the mini library generation workflow to generate a new, dataset-specific mini library was more beneficial. While identifications reduced to 350 with this “regenerated mini library”, preventing improvements from occurring at count level, the overall accuracy of the regenerated mini library increased to 91.4%, from 77.8% with the predicted library.

However, the greatest proportional improvement (TN, 60.5% vs 61.8%; TP, 30.7 vs 34.2%; FN, 4.0% vs 2.2%; FP, 4.8% vs 1.9%; overall accuracy, 77.8% vs 96.0% for predicted and shared mini library, respectively) was achieved using a third mini library which was restricted to peptides with identifications shared by both the *in silico*-predicted and regenerated mini library, though this came at the cost of further reduced identifications – 322 from 397 total identified proteins (Supplementary Figure 4).

## 4 Discussion

As improvements in DIA-MS technology continue, with instruments and software delivering improved speed, sensitivity and proteome coverage, there is an ongoing need to improve the quality of the quantification of the growing number of identified proteins. Multiple studies regarding spectral library optimisation have focused on methods for increasing proteome coverage; fewer have focussed primarily on improving resulting quantification accuracy or precision (Ge et al., 2021; Wen et al., 2025a; Rice and Belani, 2022; Wang et al., 2022). Additionally, the validation of optimised MS workflows, whether that be changes to practical methods, spectral library generation or analysis software, often emphasises the importance of %CV as a key metric for determining success, however, lower or equal %CV does not always translate to greater statistical power to detect differential protein abundance (Alharbi, Campbell and Unwin, submitted for publication). In contrast, our method prioritises improving quantification accuracy and precision of known abundance ratios, and the capacity to discriminate between differentially abundant proteins, rather than maximising the number of protein identifications and reducing the %CV.

Not all peptides demonstrate equal quantitative performance, for example, low abundance peptides with low signal-to-noise ratios present quantitative challenges, as do the inherent properties of certain peptides which leads to non-uniform protein representation such as those with chimeric spectra, post-translational modifications, and the effects of protein structure on enzymatic kinetics and peptide stability, as well as the effects of matrix interference (Parker, Venkatraman and Van Eyk, 2016; Shteynberg et al., 2016; Dorfer et al., 2018; Dupree et al., 2020; Fang et al., 2020, Chambers et al., 2007). Spectral library composition has been shown to have a significant impact on quantification and downstream differential abundance analysis, and the considerations for identification accuracy and quantification accuracy are intrinsically different (Parker, Venkatraman and Van Eyk, 2016). Thus, we have shown that an approach to spectral library optimisation in which the quality and reliability of peptides incorporated is central, rather than focussing on the quantity of peptides in the spectral library or identified during analysis, can result in a more accurate and precise representation of relative protein abundance.

Here we demonstrate that removing precursors from the spectral library used for data analysis which fail to meet specific quality criteria for quantification (creating a so-called “mini library”) can lead to minimal losses in protein identifications whilst substantially improving protein quantification performance across multiple benchmarks and reducing computational demands.

Through iterative testing of mini library filters, we developed a workflow which filters by CV and number of missing values and either expected ratio or coefficient of determination. When comparing filtering methods, we prioritised balance between improved true positive and reduced false identifications whilst retaining adequate identification counts. Many filter combinations, whilst superior in terms of proportional binary classification metrics, resulted in excessive identification loss – up to and over 50% – rendering them impractical. This is unsurprising given the well-established observation that larger spectral libraries yield increased peptide and protein identifications, thus, it is reasonable that filtering precursors from the spectral library would inevitably lead to some identification loss (Parker, Venkatraman and Van Eyk, 2016).

However, our final filtering criteria achieved a substantial reduction in library size when compared to minimal identification loss, removing nearly 5 million precursors (unidentified or poorly quantitative) and losing only 12% of protein identifications. Of precursors present in the samples, approximately 34% were deemed poor quality, and unsuitable for inclusion in the spectral library. Lost protein identifications were predominantly of poor quality, characterized by single or absent proteotypic peptides and extensive missing values – features that compromise both identification confidence and quantification reliability measurements (Carr et al., 2004; Bramer et al., 2020; Harris et al., 2023). As such these proteins would, or should, be discounted by downstream filtering steps to determine differential intensity in any case. Reducing the number of spectral library precursors by such an extent, speeds up processing time whilst limiting analysis to high-quality precursors. The balance we achieved between spectral library size and protein identifications is particularly valuable given that the field has historically used the number of identifications and DAPs as benchmarks of performance. There is an understandable reluctance to sacrifice a substantial number of identifications. However, our approach represents a deliberate compromise: modest protein identification loss in exchange for enhanced quantification quality and more reliable classification of proteins with differential abundance.

To justify this approach, we demonstrated that the mini library could achieve improved quantitative performance. The mini library consistently outperformed both comparison libraries – predicted via *in silico* processing of a species-specific sequence database and an empirical library built from all precursors identified in the sample – based on the binary classification of proteins across multiple analytical approaches. Crucially, these statistical improvements translated to actual enhancements in quantification precision as shown by LFQbench analysis, improved protein abundance ratio accuracy and improved differential abundance analysis – particularly for differentially abundant *E. coli* proteins in our model system – suggesting that our filtering criteria effectively distinguished between high- and low-quality precursors whose unreliability negatively impacts protein-level quantifications, degrading overall data quality, and demonstrating the practical value of mini library use in typical experimental workflows.

Whilst improvements with the mini library are clear, we acknowledge that the filtering criteria were tailored to this particular dataset. Transferability of the original mini library to data acquired using a substantially different LC-MS method — with a halved gradient and 10-fold reduction in target analyte levels — was expectedly limited. Under such conditions, the matrix background and co-eluting peptide complement will differ markedly, altering the interference landscape such that transitions performing well in one method may perform poorly in another, and vice versa. This is not a limitation of the mini library concept, but rather a reflection of the method-dependency inherent to transition selection.

Critically, we did replicate the approach in a second dataset using the same filtering parameters without modification, confirming that generation of a method-specific mini library remained beneficial, although bespoke adjustment of filtering parameters may be necessary to optimize performance. This underscores that the mini library approach is best understood as a fixed-method tool: established once within a defined analytical workflow and then deployed across the large sample cohorts or repeated analyses for which it is designed, delivering gains in quantitative accuracy and computational efficiency. For such cohort studies, where a consistent LC-MS method is a practical prerequisite, or settings with established workflows and sample types, the method-specificity of the mini library is a feature rather than a limitation.

Notably, merging the mini libraries generated from two distinct analytical methods did generate a smaller yet consistent set of peptides which produced further enhancements in quantitative accuracy despite a significant lowering of total identifications. While this is unsuitable for large-scale DIA studies, these peptides likely serve as strong candidates for downstream targeted MS method development at either the single protein or larger scale, particularly given recent advances in highly multiplexed targeted platforms (Plubell et al., 2026).

This work therefore serves primarily as a proof of principle, demonstrating the viability of the mini library approach whilst recognising that, currently, the library needs to be built around a specific LC-MS workflow. Future work combining and determining an intersect of mini libraries from different, deeper coverage datasets looking for consistently accurate and reliable peptides across platforms and experimental workflows may begin to enable development of curated mini libraries for common sample types, such as plasma, which could enable improved quantification quality without requiring individual laboratories to perform time- and computationally intensive library generation and filtering. However, this requires much deeper investigation.

We also acknowledge that the mini libraries generated here demonstrate improved detection of differentially abundant *E. coli* proteins since *E. coli* proteins were spiked at defined concentrations, providing clearer benchmarks for assessing quantification performance. While human peptides in the library are selected to be reproducible, they do not scale so linearity of signal cannot be guaranteed. Thus, to build a mini library for the quantification of human plasma proteins, future work should incorporate human protein spike-ins across a broader concentration range into a relevant background to better evaluate improvements in quantifying endogenous human proteins, which is the ultimate goal of this approach. Furthermore, the integration of additional quality metrics beyond those explored here – such as chromatographic peak shape, or fragment ion selection as demonstrated by Vu et al., (2026) – might enable even more refined selection of high-quality peptides.

In summary, here we challenge the common assumption that maximizing identifications is the primary, and most consequential goal of proteomics method development. We would argue that our method – which produces fewer identifications, but higher quality quantifications – provides more value than a larger dataset impaired by unreliable measurements. Our results suggest that strategic library refinement can shift the balance from maximizing identifications towards optimizing quantification quality – a compromise which may be particularly valuable in targeted, diagnostic and other clinical studies where accurate quantification and identification of differentially abundant proteins is paramount. The reduced false positive rate is particularly beneficial. False discoveries can lead to misallocation of resources in follow-up validation studies, incorrect biological interpretations, obscurement of valuable results within noise or reduction of the statistical power of subsequent analyses including interaction networks and gene ontology analysis (Wen et al., 2025b). Likewise, decreasing false negatives reduces the likelihood that valuable candidate biomarkers of biologically-relevant changes are missed purely due to a subset of poorly representative peptides.

Overall, we believe that mini library generation and the resulting improvements in DIA quantification accuracy and precision represent a meaningful step forward in quantitative proteomics.

## 5 Associated Data

The mass spectrometry proteomics data have been deposited to the ProteomeXchange Consortium via the PRIDE partner repository with the dataset identifier PXD075041. All code is available upon request from the study authors.

## Acknowledgements

BAB was funded by The Macular Society UK. HA was funded by Taibah University, Saudi Arabia. Mass spectrometry analysis was performed in part at the Stoller Biomarker Discovery Centre which was funded through the UK Medical Research Council (MR/M008959/1) and was facilitated by the Manchester National Institute for Health Research Biomedical Research Centre and the Greater Manchester Comprehensive Local Research Network. We would like to acknowledge the support of the Biomolecular Analysis Core Facility at The University of Manchester for the gift of *E. coli* pellets, and the Biological Mass Spectrometry Core Facility, part of the Stoller Clinical Analysis and Research Facility (SCARF) at the University of Manchester, in particular Drs David Knight, Stacey Warwood, and George Taylor for support with MS analysis (RRID code: SCR_020987).

The authors have declared no conflict of interest

## Abbreviations

DIA: Data Independent Acquisition
MBR: Match Between Runs
CV: Coefficient of Variation
AUC: Area Under Curve
ROC: Receiver Operator Curve
DAA: Differential Abundance Analysis
FPR: False Positive Rate
DAP: Differentially Abundant Protein
TP: True Positive
TN: True Negative
FP: False Positive
FN: False Negative

