## Supplementary figures and tables for "Enhanced proteome relative quantification using refined quantotypic spectral libraries"

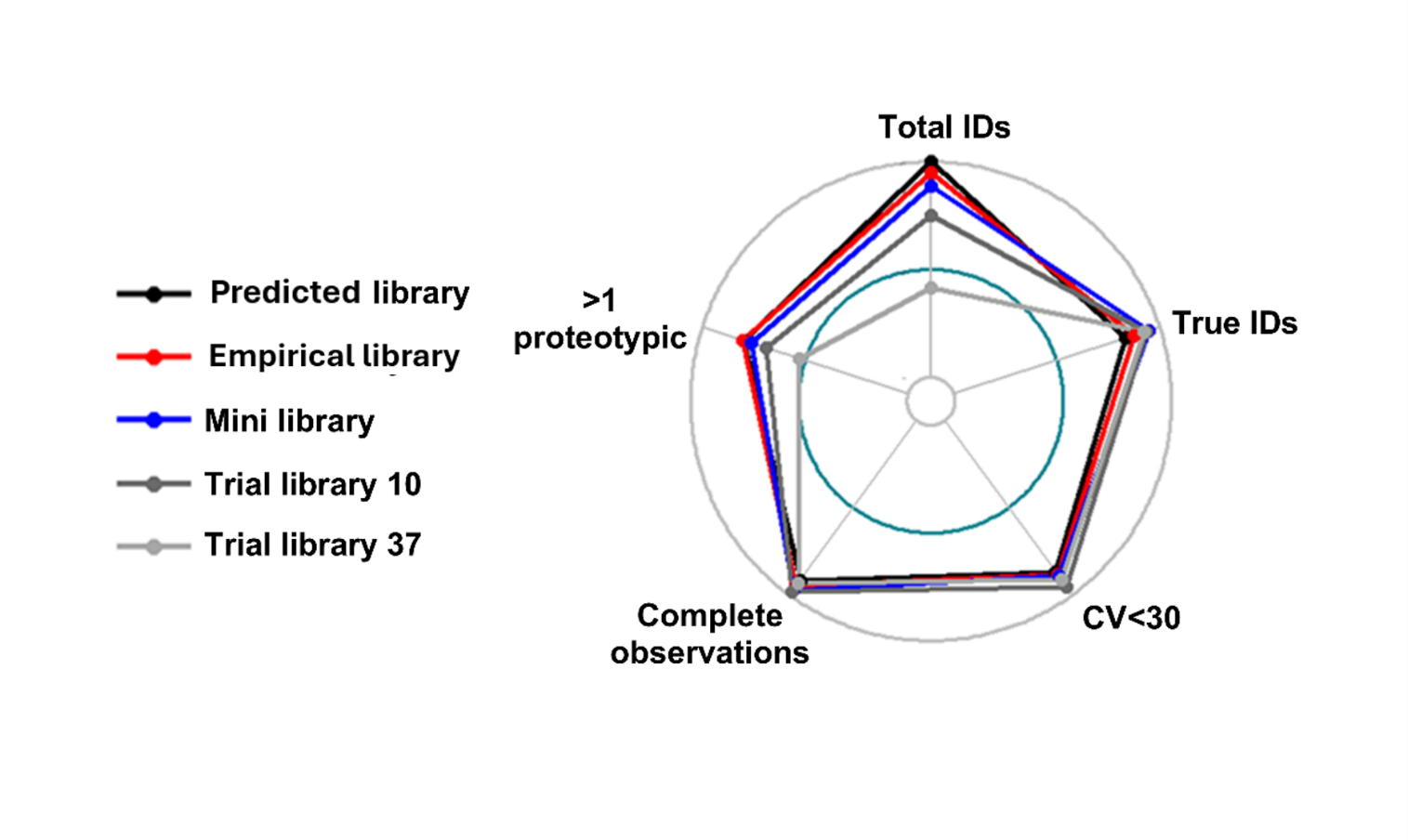
**Supplementary Figure 1. Radar plot of spectral libraries and their performance**

Quality metrics considered when assessing the performance of the predicted, empirical, chosen mini library and two example trial mini libraries with alternative quality filter metrics. Metrics considered were the percentage of protein identifications in each library compared to the 797 proteins identified with the predicted library (Total IDs), the percentage of true positive plus true negative identifications compared to the total number of protein identifications for the associated library (True IDs), the percentage of proteins with coefficient of variation less than 30% (CV <30), the percentage of proteins with identifications in all technical replicates (Complete observations) and the percentage of proteins which have 1 or more proteotypic peptides from the total number of identified proteins from the corresponding library (>1 proteotypic).

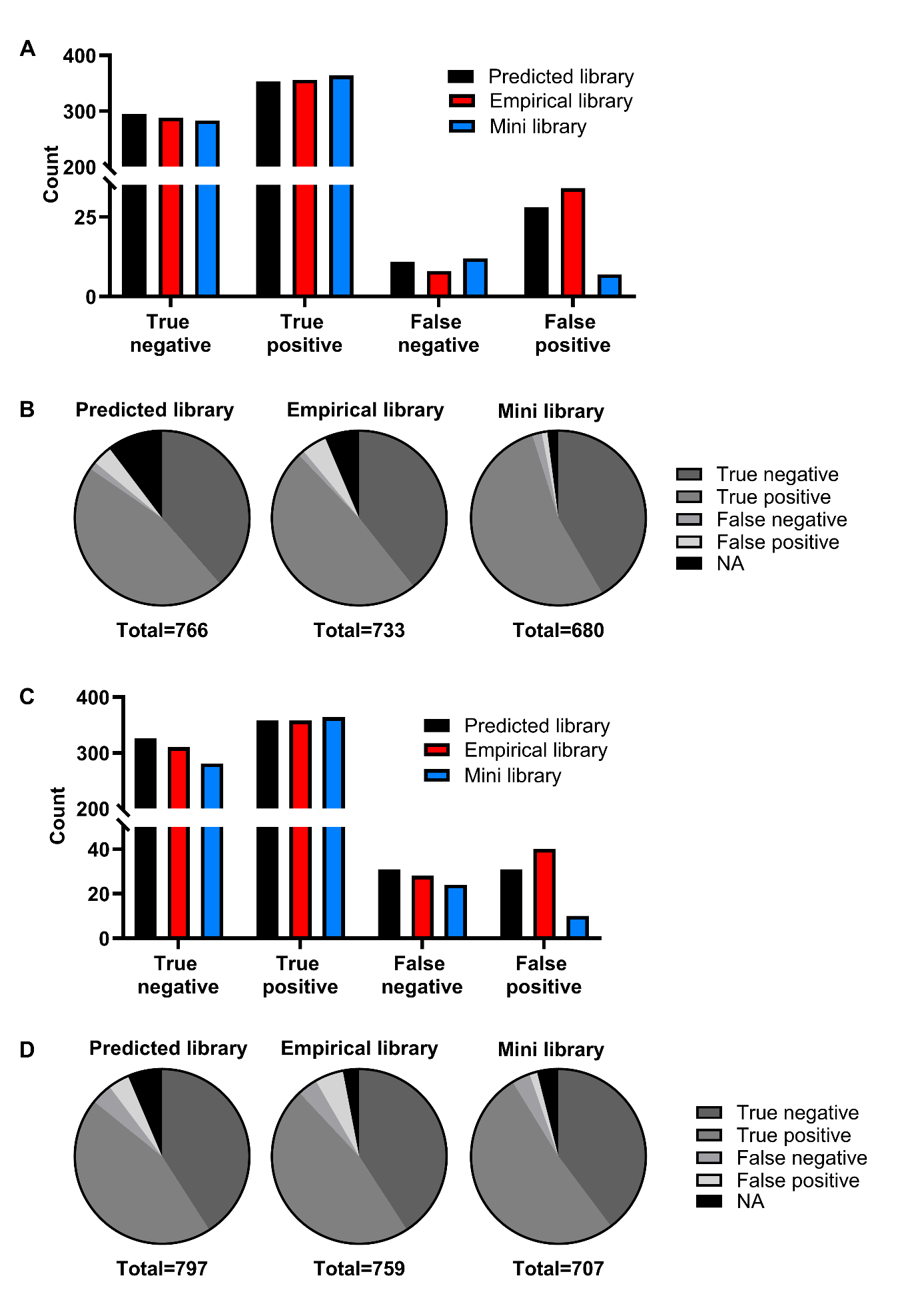
**Supplementary Figure 2. Mini library improves accuracy of binary classification across multiple analytical approaches**

An unpaired Student's t-test was performed between E4 and E7.5 protein quantifications for human and *E. coli* identifications to determine true positives (TP; significantly changed *E. coli* proteins), true negatives (TN; stable human proteins), false positives (FP; falsely significant human proteins), and false negatives (FN; non-significant *E. coli* proteins). (A) Raw counts of TP, TN, FP, and FN classifications for each library using mean imputation and excluding identifications with one or fewer proteotypic peptides. (B) Proportions of TP, TN, FP, FN, and NA (excluded identifications) categories from panel A for each library. (C) Raw counts of TP, TN, FP and FN classifications for each library with unequal sample sizes. (D) Proportions of TP, TN, FP, FN, and NA categories from panel C for each library.

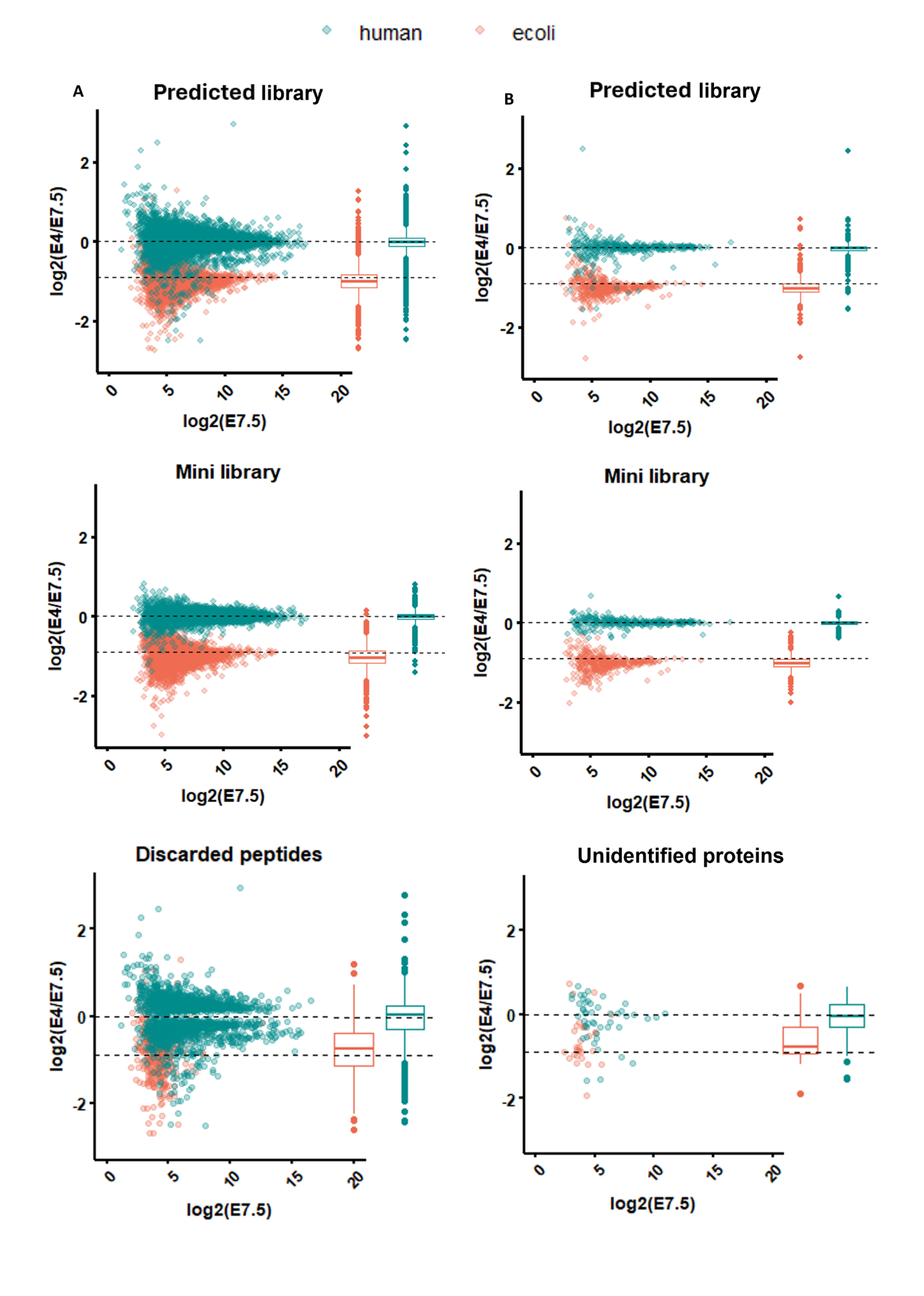
**Supplementary Figure 3. Improved precision at the peptide and protein level**

LFQbench-type analysis of the performance of the predicted and mini libraries at (A) peptide level and (B) protein level. Discarded peptides are peptides that were identified in the samples but were excluded from inclusion in the mini library. Unidentified proteins are those which were identified with the predicted library, but not with the mini library.

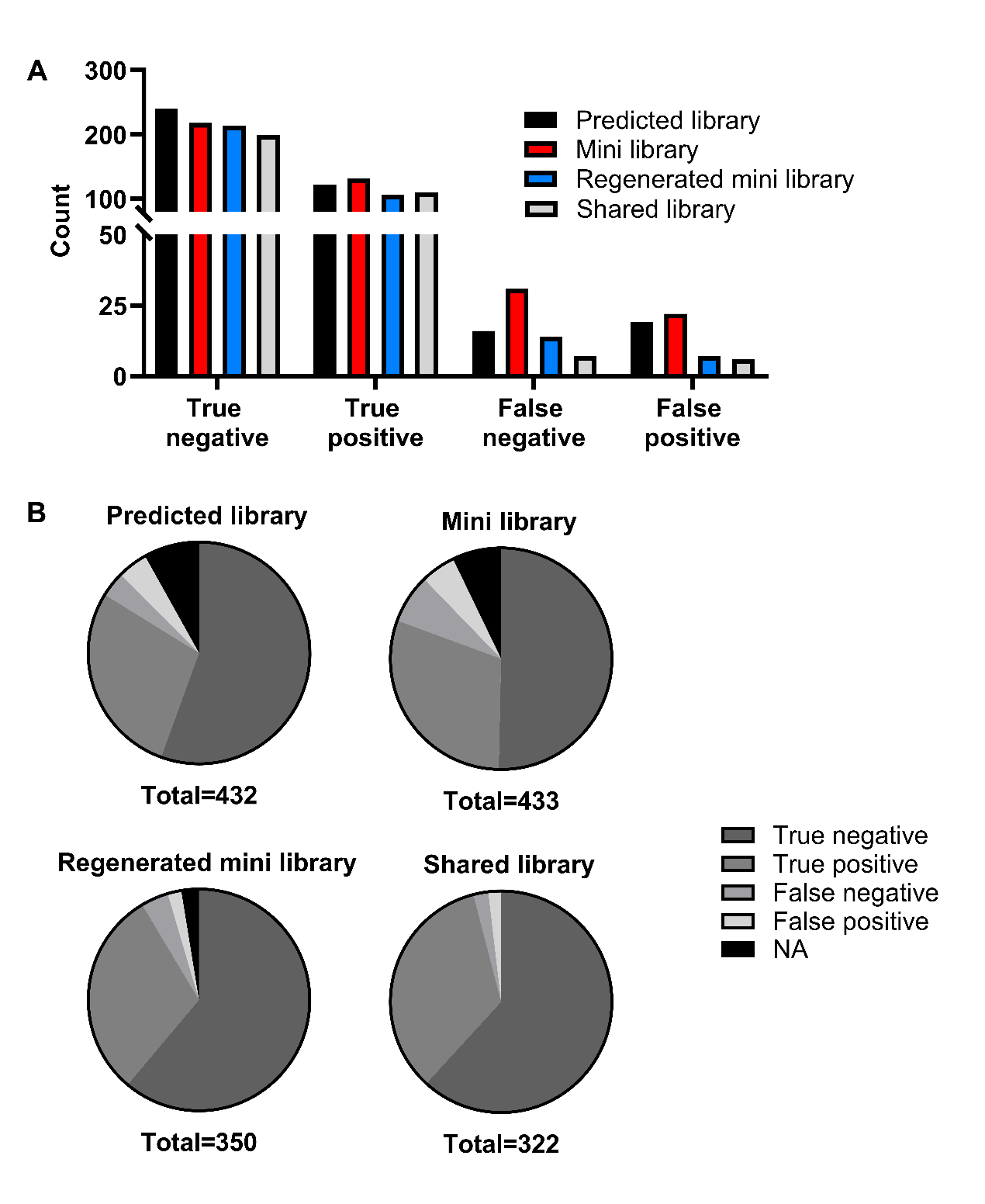

**Supplementary Figure 4. Applicability of mini libraries to an alternative dataset**

An unpaired Student's t-test was performed between 0.6ng and 1.2ng *E. coli* spike replicates from a second mixed-species dataset when analysed with the *in silico* predicted library, the mini library produced from the first dataset, a new mini library generated by applying the same filtering criteria to the second dataset (Regenerated mini library) and a library containing only precursors which were shared between the mini and regenerated libraries (Shared library), to determine true positives, true negatives, false positives and false negatives. (A) Raw counts of classifications for each library using mean imputation for proteins with fewer than three missing values; proteins with three or more missing values were excluded. (B) Proportions of classifications and NA (excluded identifications) categories from panel A.

**Supplementary Table 1. List of tested mini library filters**

| Library | Description |
| --- | --- |
| Trial lib 1 | Human: all CV <30 & all ratio +/- 30%  E. coli: 4/6 of CV <30 & ratio +/- 30% |
| Trial lib 2 | Human: all CV <20 & all ratio +/- 20%  E. coli: 4/6 of CV <20 & all ratio +/- 20% |
| Trial lib 3 | Human: all CV <20 & all ratio +/- 30%  E. coli: 6/6 of CV <20 & all ratio +/- 30% |
| Trial lib 4 | Human: all CV <20 & all ratio +/- 20%  E. coli: 6/6 of CV <20 & all ratio +/- 20% |
| Trial lib 5 | Human: all CV <20 & all ratio +/- 30%  E. coli: 4/6 of CV <20 & all ratio +/- 30% |
| Trial lib 6 | Human: all CV <30 & all ratio +/- 30%  E. coli: 6/6 of CV <30 and ratio +/- 30% |
| Trial lib 7 | Human: all CV <30 & all ratio +/- 20%  E. coli: 4/6 of CV <30 & ratio +/- 20% |
| Trial lib 8 | Human: all CV <30 & all ratio +/- 20%  E. coli: 6/6 of CV <30 & ratio +/- 20% |
| Trial lib 9 | Human: all CV <35 & all ratio +/- 35%  E. coli: 4/6 of CV <35 and ratio +/- 35% |
| Trial lib 10 | Human: all CV <35 & all ratio +/- 35%  E. coli: 6/6 of CV <35 and ratio +/- 35% |
| Trial lib 11 | Human: all CV <30 & all ratio +/- 30%  E. coli: 2/3 of CV <30 and 2/3 ratio +/- 30% |
| Trial lib 12 | Human: all CV <20 & all ratio +/- 20%  E. coli: 2/3 of CV <20 & 2/3 ratio +/- 20% |
| Trial lib 13 | Human: all CV <20  E. coli: all CV <20 |
| Trial lib 14 | Human: all CV <30  E. coli: all CV <30 |
| Trial lib 15 | Human: all ratio +/- 20%  E. coli: all ratio +/- 20% |
| Trial lib 16 | Human: all ratio +/- 30%  E. coli: all ratio +/- 30% |
| Trial lib 17 | Human: all ratio +/- 15%  E. coli: all ratio +/- 15% |
| Trial lib 18 | Human: all ratio +/- 0.1  E. coli: all ratio +/- 0.1 |
| Trial lib 19 | Human: all ratio +/- 0.2  E. coli: all ratio +/- 0.2 |
| Trial lib 20 | Human: 2/3 of CV <30 & R^2^ <0.25 OR 2/3 of CV <30 & all ratio +/- 25%  E. coli: 2/3 of CV <30 & R^2^ >0.75 & R >0 OR 2/3 of CV <30 & all ratio +/- 15% |
| Trial lib 21 | Human: 2/3 of CV <30 & R^2^ <0.25 OR 2/3 of CV <30 & all ratio +/- 15%  E. coli: 2/3 of CV <30 & R^2^ >0.75 & R >0 OR 2/3 of CV <30 & all ratio +/- 15% |
| Trial lib 22 | Human: 2/3 of CV <30 & R^2^ <0.25 OR 2/3 of CV <30 & all ratio +/- 12.5%  E. coli: 2/3 of CV <30 & R^2^ >0.75 & R >0 OR 2/3 of CV <30 & all ratio +/- 12.5% |
| Trial lib 23 | Human: 2/3 of CV <30 & R^2^ <0.25 OR 2/3 of CV <30 & all ratio +/- 10%  E. coli: 2/3 of CV <30 & R^2^ >0.75 & R >0 OR 2/3 of CV <30 & all ratio +/- 10% |
| Trial lib 24 | Human: 2/3 of CV <30 & R^2^ <0.25 OR 2/3 of CV <30 & all ratio +/- 0.1  E. coli: 2/3 of CV <30 & R^2^ >0.75 & R >0 OR 2/3 of CV <30 & all ratio +/- 0.1 |
| Trial lib 25 | Human: 2/3 of CV <30 & R^2^ <0.25 OR 2/3 of CV <30 & all ratio +/- 0.2  E. coli: 2/3 of CV <30 & R^2^ >0.75 & R >0 OR 2/3 of CV <30 & all ratio +/- 0.2 |
| Trial lib 26 | Human: all CV <30 & R^2^ <0.25 OR 2/3 of CV <30 & all ratio +/- 15%  E. coli: all CV <30 & R^2^ >0.75 & R >0 OR 2/3 of CV <30 & all ratio +/- 15% |
| Trial lib 27 | Human: mean CV <30 & R^2^ <0.25 OR 2/3 of CV <30 & all ratio +/- 15%  E. coli: mean CV <30 & R^2^ >0.75 & R >0 OR 2/3 of CV <30 & all ratio +/- 15% |
| Trial lib 28 | Human: 2/3 of CV <30 & R^2^ <0.3 OR 2/3 of CV <30 & all ratio +/- 25%  E. coli: 2/3 of CV <30 & R^2^ >0.8 & R >0 OR 2/3 of CV <30 & all ratio +/- 15% |
| Trial lib 29 | Human: 2/3 of CV <30 & R^2^ <0.3 OR 2/3 of CV <30 & all ratio +/- 15%  E. coli: 2/3 of CV <30 & R^2^ >0.8 & R >0 OR 2/3 of CV <30 & all ratio +/- 15% |
| Trial lib 30 | Human: 2/3 of CV <30 & R^2^ <0.3 OR 2/3 of CV <30 & all ratio +/- 12.5%  E. coli: 2/3 of CV <30 & R^2^ >0.8 & R >0 OR 2/3 of CV <30 & all ratio +/- 12.5% |
| Trial lib 31 | Human: 2/3 of CV <30 & R^2^ <0.3 OR 2/3 of CV <30 & all ratio +/- 10%  E. coli: 2/3 of CV <30 & R^2^ >0.8 & R >0 OR 2/3 of CV <30 & all ratio +/- 10% |
| Trial lib 32 | Human: 2/3 of CV <30 & R^2^ <0.3 OR 2/3 of CV <30 & all ratio +/- 0.1  E. coli: 2/3 of CV <30 & R^2^ >0.8 & R >0 OR 2/3 of CV <30 & all ratio +/- 0.1 |
| Trial lib 33 | Human: 2/3 of CV <30 & R^2^ <0.3 OR 2/3 of CV <30 & all ratio +/- 0.2  E. coli: 2/3 of CV <30 & R^2^ >0.8 & R >0 OR 2/3 of CV <30 & all ratio +/- 0.2 |
| Trial lib 34 | Human: all CV <30 & R^2^ <0.3 OR 2/3 of CV <30 & all ratio +/- 15%  E. coli: all CV <30 & R^2^ >0.8 & R >0 OR 2/3 of CV <30 & all ratio +/- 15% |
| Trial lib 35 | Human: mean CV <30 & R^2^ <0.3 OR 2/3 of CV <30 & all ratio +/- 15%  E. coli: mean CV <30 & R^2^ >0.8 & R >0 OR 2/3 of CV <30 & all ratio +/- 15% |
| Trial lib 36 | Human: R^2^ <0.3 OR 2/3 of CV <30 & all ratio +/- 15%  E. coli: R^2^ >0.8 & R >0 OR 2/3 of CV <30 & all ratio +/- 15% |
| Trial lib 37 | Human: R^2^ <0.3 & all ratio +/- 15%  E. coli: R^2^ >0.8 & all ratio +/- 15% |
| Trial lib 38 | Human: R^2^ <0.3 OR 2/3 of CV <30 & all ratio +/- 0.1  E. coli: R^2^ >0.8 & R >0 OR 2/3 of CV <30 & all ratio +/- 0.1 |
| Trial lib 39 | Human: 2/3 of CV <30 & R^2^ <0.3 OR 2/3 of CV <30 & all ratio +/- 25%  E. coli: 2/3 of CV <30 & R^2^ >0.7 & R >0 OR 2/3 of CV <30 & all ratio +/- 15% |
| Trial lib 40 | Human: 2/3 of CV <30 & R^2^ <0.3 OR 2/3 of CV <30 & all ratio +/- 15%  E. coli: 2/3 of CV <30 & R^2^ >0.7 & R >0 OR 2/3 of CV <30 & all ratio +/- 15% |
| Trial lib 41 | Human: 2/3 of CV <30 & R^2^ <0.3 OR 2/3 of CV <30 & all ratio +/- 12.5%  E. coli: 2/3 of CV <30 & R^2^ >0.7 & R >0 OR 2/3 of CV <30 & all ratio +/- 12.5% |
| Trial lib 42 | Human: 2/3 of CV <30 & R^2^ <0.3 OR 2/3 of CV <30 & all ratio +/- 10%  E. coli: 2/3 of CV <30 & R^2^ >0.7 & R >0 OR 2/3 of CV <30 & all ratio +/- 10% |
| Trial lib 43 | Human: 2/3 of CV <30 & R^2^ <0.3 OR 2/3 of CV <30 & all ratio +/- 0.1  E. coli: 2/3 of CV <30 & R^2^ >0.7 & R >0 OR 2/3 of CV <30 & all ratio +/- 0.1 |
| Trial lib 44 | Human: 2/3 of CV <30 & R^2^ <0.3 OR 2/3 of CV <30 & all ratio +/- 0.2  E. coli: 2/3 of CV <30 & R^2^ >0.7 & R >0 OR 2/3 of CV <30 & all ratio +/- 0.2 |
| Trial lib 45 | Human: all CV <30 & R^2^ <0.3 OR 2/3 of CV <30 & all ratio +/- 15%  E. coli: all CV <30 & R^2^ >0.7 & R >0 OR 2/3 of CV <30 & all ratio +/- 15% |
| Trial lib 46 | Human: mean CV <30 & R^2^ <0.3 OR 2/3 of CV <30 & all ratio +/- 15%  E. coli: mean CV <30 & R^2^ >0.7 & R >0 OR 2/3 of CV <30 & all ratio +/- 15% |
| Trial lib 47 | Human: 2/3 of CV <30 & R^2^ <0.4 OR 2/3 of CV <30 & all ratio +/- 25%  E. coli: 2/3 of CV <30 & R^2^ >0.6 & R >0 OR 2/3 of CV <30 & all ratio +/- 15% |
| Trial lib 48 | Human: 2/3 of CV <30 & R^2^ <0.4 OR 2/3 of CV <30 & all ratio +/- 15%  E. coli: 2/3 of CV <30 & R^2^ >0.6 & R >0 OR 2/3 of CV <30 & all ratio +/- 15% |
| Trial lib 49 | Human: 2/3 of CV <30 & R^2^ <0.4 OR 2/3 of CV <30 & all ratio +/- 12.5%  E. coli: 2/3 of CV <30 & R^2^ >0.6 & R >0 OR 2/3 of CV <30 & all ratio +/- 12.5% |
| Trial lib 50 | Human: 2/3 of CV <30 & R^2^ <0.4 OR 2/3 of CV <30 & all ratio +/- 10%  E. coli: 2/3 of CV <30 & R^2^ >0.6 & R >0 OR 2/3 of CV <30 & all ratio +/- 10% |
| Trial lib 51 | Human: 2/3 of CV <30 & R^2^ <0.4 OR 2/3 of CV <30 & all ratio +/- 0.1  E. coli: 2/3 of CV <30 & R^2^ >0.6 & R >0 OR 2/3 of CV <30 & all ratio +/- 0.1 |
| Trial lib 52 | Human: 2/3 of CV <30 & R^2^ <0.4 OR 2/3 of CV <30 & all ratio +/- 0.2  E. coli: 2/3 of CV <30 & R^2^ >0.6 & R >0 OR 2/3 of CV <30 & all ratio +/- 0.2 |
| Trial lib 53 | Human: all CV <30 & R^2^ <0.4 OR 2/3 of CV <30 & all ratio +/- 15%  E. coli: all CV <30 & R^2^ >0.6 & R >0 OR 2/3 of CV <30 & all ratio +/- 15% |
| Trial lib 54 | Human: mean CV <30 & R^2^ <0.4 OR 2/3 of CV <30 & all ratio +/- 15%  E. coli: mean CV <30 & R^2^ >0.6 & R >0 OR 2/3 of CV <30 & all ratio +/- 15% |

**Supplementary Table 2. Percentage of classifications based upon t-tests under multiple analytical conditions for each library with the superior result between the libraries in green.**

| **Library** | **T-test for complete cases**  **(%)** | | | | **T-test after mean imputation of ≤2 missing values**  **(%)** | | | | **T-test after mean imputation and single/ no proteotypic peptide ID removal**  **(%)** | | | | **T-test with unequal sample sizes**  **(%)** | | | |
| --- | --- | --- | --- | --- | --- | --- | --- | --- | --- | --- | --- | --- | --- | --- | --- | --- |
|  | **TN** | **TP** | **FN** | **FP** | **TN** | **TP** | **FN** | **FP** | **TN** | **TP** | **FN** | **FP** | **TN** | **TP** | **FN** | **FP** |
| **Predicted** | 35.5 | 35.3 | 0.25 | 3.3 | 39.3 | 44.4 | 1.4 | 4.1 | 38.2 | 45.7 | 1.4 | 3.6 | **40.9** | 44.9 | 3.9 | 3.9 |
| **Empirical** | 37 | 38.6 | **0.1** | 4.3 | 40.7 | 47 | **1.1** | 4.9 | 39.2 | 48.5 | **1.1** | 4.6 | 40.8 | 47.2 | 3.7 | 5.3 |
| **Mini** | **38.9** | **44.3** | 0.7 | **1.3** | **43.1** | **51.6** | 1.7 | **1.4** | **41.6** | **53.5** | 1.8 | **1** | 39.7 | **51.5** | **3.4** | **1.4** |

**Supplementary Table 3. LFQbench accuracy and precision metrics**

| Ratio comparison | Human | | *E. coli* | |
| --- | --- | --- | --- | --- |
|  | **Predicted** | **Mini** | **Predicted** | **Mini** |
| Protein global precision | 0.21 | 0.17 | 0.26 | 0.21 |
| Protein global accuracy | 0.00 | 0.00 | -0.09 | -0.09 |
| Protein technical variance | 0.08 | 0.06 | - | - |
| Peptide global precision | 0.30 | 0.13 | 0.34 | 0.27 |
| Peptide global accuracy | 0.00 | 0.00 | -0.08 | -0.10 |
| Peptide technical variance | 0.13 | 0.10 | - | - |

**Supplementary Table 4. LFQbench global species overlap metrics**

| Ratio comparison | Predicted | Mini |
| --- | --- | --- |
| Protein global species overlap | 0.885 | 0.999 |
| Peptide global species overlap | 0.973 | 0.998 |

**Supplementary Table 5. Mini library improves the count of identification ratios which lie within +/- 20% of the expected abundance ratio for proteins identified in both libraries.**

| Ratio comparison | Human | | *E. coli* | | Total | | |
| --- | --- | --- | --- | --- | --- | --- | --- |
|  | **Predicted** | **Mini** | **Predicted** | **Mini** | **Predicted** | | **Mini** |
| E7.5 vs E4 | 295 | 297 | 306 | 310 | 601 | 607 | |
| E7.5 vs E5 | 291 | 284 | 317 | 330 | 608 | 614 | |
| E5 vs E4 | 286 | 275 | 333 | 339 | 619 | 614 | |

**Supplementary Table 6. Number of differentially abundant and stable proteins for each pairwise concentration comparison.**

| Ratio comparison | Library | Differentially abundant | | Stably abundant | |
| --- | --- | --- | --- | --- | --- |
|  |  | **True** | **False** | **True** | **False** |
| E7.5 vs E4 | **Predicted** | 341 | 1 | 368 | 87 |
|  | **Mini** | 345 | 0 | 317 | 45 |
| E7.5 vs E5 | **Predicted** | 319 | 0 | 369 | 109 |
|  | **Mini** | 312 | 0 | 317 | 78 |
| E5 vs E4 | **Predicted** | 137 | 1 | 368 | 291 |
|  | **Mini** | 156 | 2 | 315 | 234 |
